# Pseudopaline, a staphylopine-like metallophore involved in zinc and nickel uptake in *Pseudomonas aeruginosa*

**DOI:** 10.1101/165571

**Authors:** Sébastien Lhospice, Nicolas Oswaldo Gomez, Laurent Ouerdane, Catherine Brutesco, Ghassan Ghssein, Christine Hajjar, Ahmed Liratni, Shuanglong Wang, Pierre Richaud, Sophie Bleves, Geneviève Ball, Elise Borezée-Durant, Ryszard Lobinski, David Pignol, Pascal Arnoux, Romé Voulhoux

## Abstract

Metal uptake is vital for all living organisms. In metal scarce conditions, a common bacterial strategy consists in the biosynthesis of metallophores, their export in the extracellular medium and the recovery of a metal-metallophore complex through dedicated membrane transporters. Staphylopine is a recently described metallophore distantly related to plant nicotianamine that contributes to the broad-spectrum metal uptake capabilities of *Staphylococcus aureus*. Here, we characterize a four genes operon (*PA4837–PA4834*) in *Pseudomonas aeruginosa* involved in the biosynthesis and trafficking of a staphylopine-like metallophore named pseudopaline. Pseudopaline differs from staphylopine with regard to the stereochemistry of its histidine moiety associated to an alpha ketoglutarate moiety instead of pyruvate. *In vivo*, the pseudopaline operon is regulated by zinc through the Zur repressor. The metal-uptake property of the pseudopaline system appears different from that of staphylopine with a predominant effect on nickel uptake, and on zinc uptake in metal scarce conditions mimicking a chelating environment, thus reconciling the regulation of the *cnt* operon by zinc with its function as a zinc importer under metal scarce conditions.

**AUTHOR SUMMARY:** Zinc is an essential micronutrients for bacteria, particularly important at the host-pathogen interface where the host tends to sequester metals in a so called nutritional immunity framework, and the pathogenic bacterium increases its metal uptake efforts in order to keep up with its metal requirements. Here we reveal a novel metallophore, named pseudopaline, which is synthesized and exported by *Pseudomonas aeruginosa* and serves for the uptake of nickel in metal poor media, and for the uptake of zinc in metal scarce conditions that mimic the chelating environment that presumably prevails within a host.

## INTRODUCTION

Divalent metals (Mn, Fe, Co, Ni, Cu and Zn) are essential micronutrients for all life forms, and acquisition of these metals is therefore vital, particularly for bacterial pathogens in the context of host-pathogen interactions. Indeed, there is a competition between the host, which tends to sequester metals in a so called nutritional immunity framework, and the pathogenic bacterium, which increases its metal uptake efforts in order to keep up with its metal requirements (1, 2). Most pathogenic bacteria produce metallophores for metal uptake, with siderophores being the most well-characterized metallophore family (3). Siderophores are synthesized within the cell through non ribosomal peptide synthases (NRPS) or through a NRPS independent system (NIS) and then are exported in the extracellular medium where they scavenge iron. Extracellular iron siderophore complexes can be recognized and actively transported into the periplasm by TonB dependent transporters (TBDT) in Gram-negative bacteria, and usually ABC transporters in both Gram-negative and Gram-positive bacteria are used for the crossing of the cytoplasmic membrane. There are many variations on this common theme and, for example, some bacteria do not produce a specific type of siderophore although they are able to use it for iron import (4). The siderophore pathway could also prevent toxic accumulation of metals, which was particularly studied in the case of *Pseudomonas aeruginosa* (5, 6). *P. aeruginosa* synthesizes two types of siderophores with high iron affinity, pyochelin and pyoverdine, the latter being a demonstrated virulence factor (7).

Metallophores specific for the uptake of metals other than iron have also been described, such as the chalcophore methanobactin involved in copper uptake in methane-oxidizing bacteria (8, 9). Manganesophore have not been described as such, although TseM, a protein effector secreted through a Type VI secretion system, was shown to play an important role in TBDT-dependent manganese uptake in *Burkholderia thailandensis* (10). There is also indirect evidence for the existence of a nickelophore in *Escherichia coli*, although it has still to be identified (11). Free histidine could also be used as a nickelophore *in vivo* for nickel uptake in various bacteria (12, 13). Yersiniabactin, initially described as a siderophore, also exhibits zincophore properties in *Yersinia pestis* (14, 15). Coelibactin, described in *Streptomyces coelicolor*, may also represent a zincophore as it is synthesized by a NRPS under the control of Zur, a zinc responsive repressor (16).

Staphylopine is a nicotianamine-like molecule that was recently described as a metallophore with remarkable broad-spectrum specificity (17). In *Staphylococcus aureus*, staphylopine is synthesized through the action of three soluble enzymes (SaCntKLM). SaCntK transforms L-histidine in D-histidine, SaCntL transfers an aminobutyrate moiety from S-adenosylmethionine (SAM) onto D-histidine, and SaCntM reductively condensates the product of SaCntL (called xNA) with pyruvate. The staphylopine biosynthesis and trafficking pathway is responsible for zinc, copper, nickel, cobalt and iron uptake, depending on the growth conditions, and this system contributes to the virulence and fitness of *S. aureus* (17–19). The *S. aureus cnt* operon is partly conserved in *P. aeruginosa*, where homologues of the *cntL* and *cntM* genes are found, albeit with 20-30% sequence identity at the protein level. Upstream of *cntL*, a gene codes a predicted outer membrane protein belonging to the TBDT family, and downstream of *cntM*, a gene codes for a predicted inner membrane protein belonging to the EamA or DMT family (drug/metabolite transporter; Figure S1). Transcriptomic approaches revealed that this gene cluster was highly expressed in a burn wound model (20). This last gene was also identified as part of a novel siderophore pathway that appeared vital for the growth of *P. aeruginosa* in airway mucus secretion (AMS) (21). Finally, through a transcriptomic study of a Zur deficient strain, these four genes were found in the top five regulated units, although most of them were annotated as hypothetical (22).Here, we show that the four above-mentioned genes (here named *cntO*, *cntL*, *cntM* and *cntI;* see supplementary table S1 for correspondence with locus tag in PAO1, PA7 and PA14 strains of *P. aeruginosa*) are part of an operon that is regulated by zinc level through the Zur repressor. Using biochemistry and metabolomics approaches, we prove that the two biosynthetic enzymes (PaCntL and PaCntM) synthesize a novel metallophore, which we named pseudopaline, and which differs from staphylopine by the presence of a D-histidine moiety instead of L-histidine, and an α-ketoglutarate moiety instead of a pyruvate. A *cntL* mutant strain is shown to be unable to synthesize pseudopaline and is impaired in its ability to import nickel in a minimal media, supplemented or not with nickel. Under more stringent conditions where a chelator such as EDTA is added to a minimal succinate (MS) medium, a condition that presumably mimics the chelating environment prevailing within a host or in AMS, we show evidence that the *cntL* mutant strain is unable to import zinc, therefore reconciling the regulation of this operon by zinc with its function as a zinc importer functioning in metal scarce conditions.

## RESULTS AND DISCUSSION

### The *cnt* operon of *P. aeruginosa* is regulated by zinc level through the zinc-responsive regulator Zur

*In silico* analysis of the *cnt* gene cluster of *P. aeruginosa* PA14 strain indicated two overlapping open reading frames between *cntL* and *cntM* and between *cntM* and *cntI*, classically observed in operonic structures (Figure S1). Further screening of the upstream *cnt* sequence for promoter regions using Bprom software (23), revealed a σ70 promoter in the 200 base-pairs upstream from the annotated *cntO* ATG codon (Figure S1). Interestingly, a putative Zur binding box “GTTATagtATAtC” can be identified overlapping the -10 box of the predicted σ70 promoter, (22, 24). This *in silico* analysis supports an operonic organization of the four *cnt* genes and strongly suggests a transcriptional activation of this operon under zinc depletion through the Zur repressor (25, 26). In order to test this hypothesis, we performed RT-PCR experiments using as templates RNA and cDNA generated from a WT PA14 strain grown in minimal succinate (MS) medium known to contains low levels of metals, including zinc (5). The successful amplification of the four *cnt* gene transcripts (Figure S1) indeed indicated their induction when cells were grown in a MS medium. The specific amplification of the three *cnt* intergenic regions confirmed that the four *cnt* genes are co-transcribed in one single transcript and therefore constitute an operon.

To validate at the protein level the transcriptional regulation of the *cnt* genes, we followed by immunoblotting the PaCntL production under various growth conditions. In this respect, we constructed a *cntL* mutant strain producing a chromosomally encoded V5-tagged PaCntL (Δ*cntL::cntL_V5_*). In this strain, the recombinant *cntL*_V5_ gene was placed under the predicted *cnt* promoter region and inserted at the *att* site of the *P. aeruginosa* genome. In agreement with our transcriptional data, immunoblotting experiments indicated that, the recombinant PaCntL_V5_ is only produced in MS medium and not in a rich medium such as the LB medium (Figure 1A). Presumably, this is due to the low metal content of the MS medium as compared to the LB medium. We then tested whether the *cntL* transcription was subject to metal repression by checking PaCntL_V5_ production in MS medium supplemented with various concentrations of the most representative metals. Dot-blot experiments showed a specific loss of PaCntL_V5_ production in MS medium supplemented with as low as 0.1 μM of ZnSO_4_. An addition of iron, nickel or cobalt at concentrations equivalent or above the one found in LB rich medium (5) has no negative effect on PaCntL_V5_ production (Figure 1B). The hypothesis of a Zur repressor regulating the *cnt* operon was then tested through the construction of a PA14Δ*cntL::cntL_V5_ zur^-^*strain. PaCntL_V5_ was still produced in the *zur* mutant strain grown in LB or MS media supplemented with 1 μM of ZnSO_4_, conditions in which Zur normally exerts its repressor activity (Figure 1C). Taken together, these data therefore demonstrate that the *cnt* operon of *P. aeruginosa* is negatively regulated by zinc, most probably through the binding of a Zn-Zur repressor complex onto the predicted Zur binding motif identified in the σ70 promoter, thus preventing the recruitment of RNA-polymerase.

**Figure 1:**
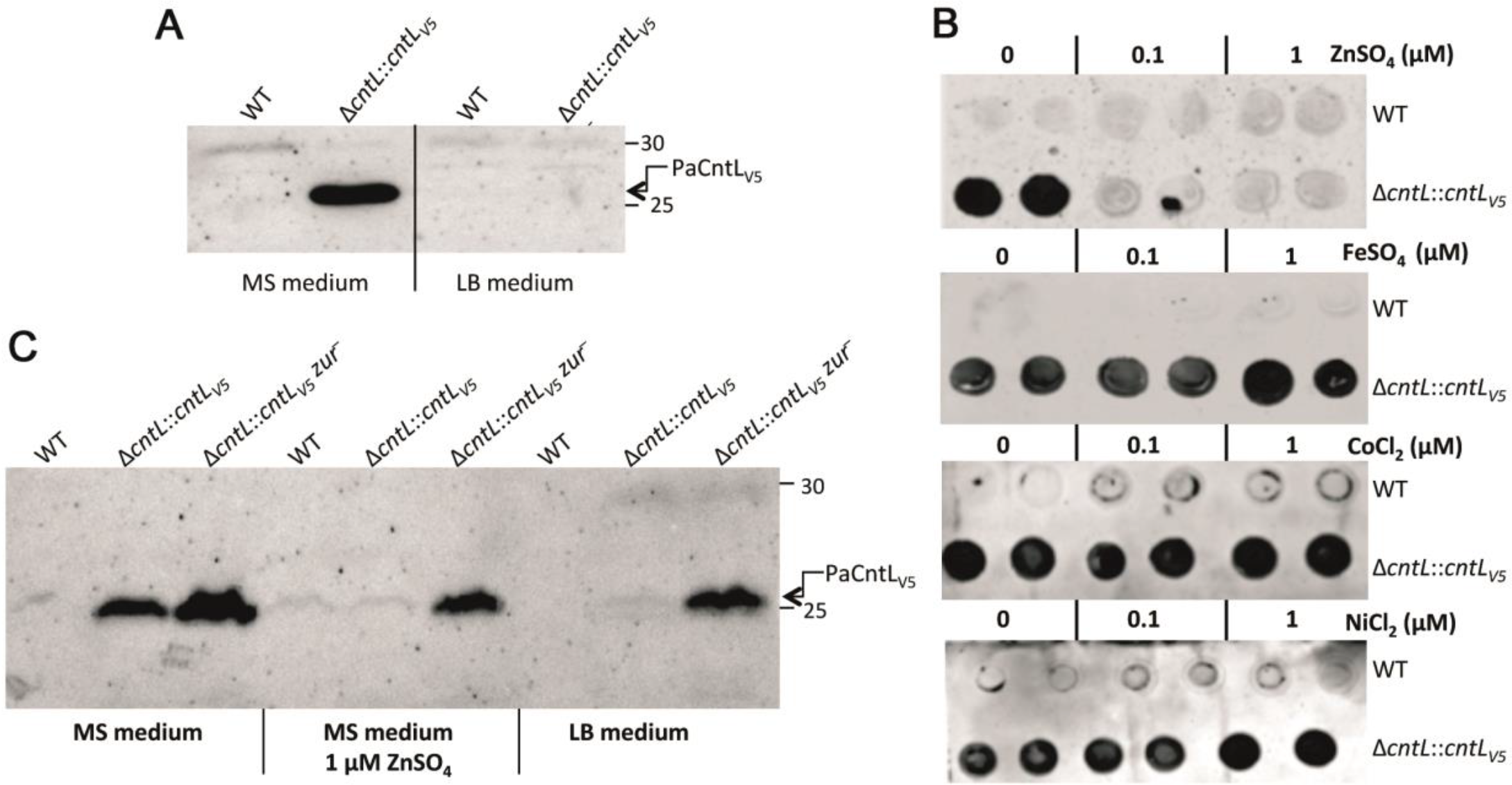
PaCntL production under various growth conditions. (A) Immunoblotting using antibody directed against the V5 epitope for revealing PaCntL_V5_ production under poor (MS) and rich (LB) media. (B) Dot-blot revealing the Pa-CntL_V5_ production in MS medium supplemented by divalent metals. (C) Immunoblot detection of PaCntL_V5_ production in PA14 WT and Zur deficient strains (*zur^-^*) in various growth conditions.

### *In vivo* detection and characterization of a PaCntL-dependent metallophore in the extracellular medium of *P. aeruginosa*

We constructed a PA14 mutant strain lacking PaCntL (Δ*cntL*) and compared the composition of the intra- and extra-cellular contents of wild type and Δ*cntL* strains grown under the previously defined *cnt* inducible conditions. Extracellular samples were analysed by hydrophilic interaction liquid chromatography (HILIC) with detection by inductively coupled plasma mass spectrometry (ICP-MS) and electrospray ionization mass spectrometry (ESI-MS). HILIC/ICP-MS data revealed the presence of a molecule complexed with nickel and zinc in the supernatant of the WT strain, which was absent in the *cntL* mutant strain (Figure 2). ESI-MS investigation of the metabolites eluting at this same elution volume showed unambiguously the presence of typical nickel and zinc isotopic patterns indicating the presence of a free metallophore with a molecular mass of 386 Da (Figure 2). Using the accurate mass and a molecular formula finder software we proposed the C_15_N_4_O_8_H_20_ empiric formula for the ligand in complex with nickel or zinc (Figure 2, inset for the nickel chelate). This ligand corresponds to a new metallophore produced by *P. aeruginosa* in a *cntL*-dependent manner. Comparison of its elemental composition with that of staphylopine (328 Da) revealed the presence of two additional carbons and two oxygen atoms, suggesting the use of an αketoglutarate(αKG) moiety instead of pyruvate as found in staphylopine. The fragmentation of this metallophore in gas-phase confirmed this hypothesis (Figure S2). We propose to name this new metallophore pseudopaline, to recall its origin from *P. aeruginosa* and its belonging to the nopaline family of opine (27).

**Figure 2:**
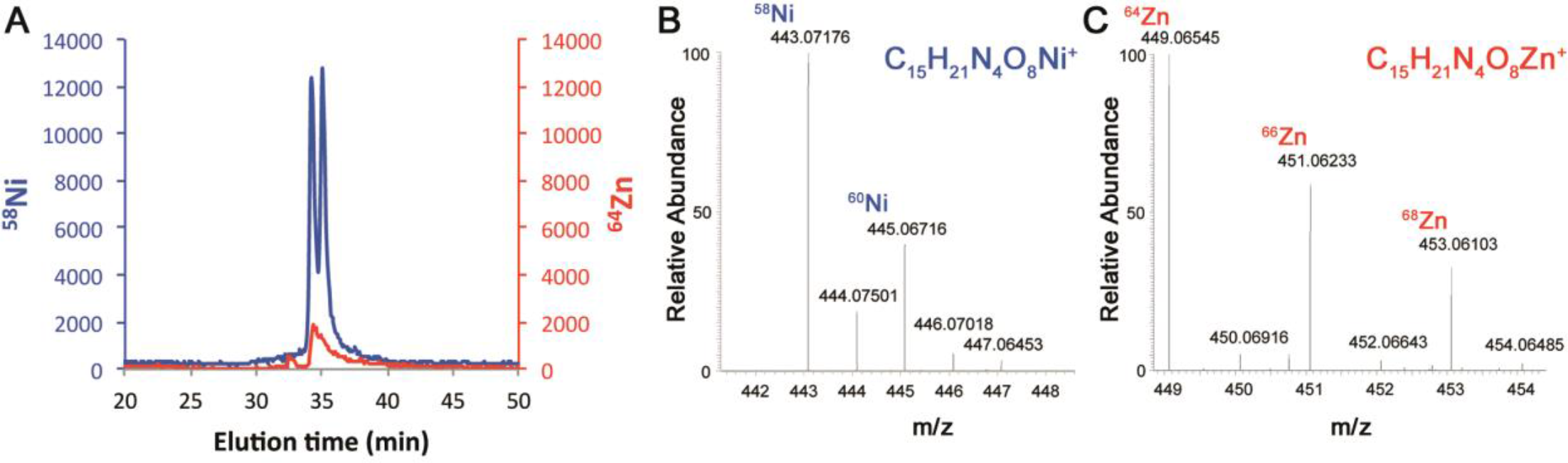
*In vivo* PaCntL-dependent detection of a nickel or zinc-bound metallophore in the extracellular fraction of *P. aeuginosa*. (A) HILIC/ICP-MS chromatogram of metal-bound metabolites. (B) HILIC-ESI/MS mass spectrum of a Ni-metallophore complex in the extracellular fraction of the WT strain but absent in the Δ*cntL* mutant. (C) HILIC-ESI/MS mass spectrum of a Zn-metallophore complex in the extracellular fraction of the WT strain but absent in the Δ*cntL* mutant. The empirical molecular formula of the CntL-dependant Ni- or Zn-metallophore complexes were deduced from the exact masse.

### *In vitro* reconstitution of the pseudopaline biosynthetic pathway catalysed by PaCntL and PaCntM

We have recently shown that the PaCntL/M orthologs in *S. aureus* (SaCntL/M) are sequentially involved in the biosynthesis of the staphylopine metallophore, using a D-histidine that is produced by the histidine racemase enzyme SaCntK (17). One of the main difference between the *cnt* operons of *P. aeruginosa* and *S. aureus* is however the absence of a *cntK* gene upstream of the *cntL-M* genes in *P. aeruginosa*. This observation led to the possibility of using directly L-histidine instead of D-histidine. In order to investigate the properties of CntL and CntM of *P. aeruginosa*, the corresponding genes were cloned, heterologously expressed in *E. coli* and their products purified for further biochemical analysis. Gel filtration experiments showed that PaCntL could form a complex with PaCntM (Figure S3), although this interaction was not observed between SaCntL and SaCntM. With regard to PaCntL, we used thin layer chromatography (TLC) separation to follow the carboxyl moiety of a carboxyl-[^14^C]-labelled S-adenosine methionine (SAM) substrate, co-incubated with either L- or D-histidine (Figure 3A). Only the incubation with L-histidine led to a novel band corresponding to a reaction intermediate that we propose to name yNA. We demonstrated subsequently that PaCntM preferentially bound to NADH and not to NADPH (Figure 3B), contrary to SaCntM that showed a preference for NADPH. We then used TLC to visualize the PaCntLM reaction products under various *in vitro* conditions using all the putative substrates (Figure 3C). Unexpectedly, the co-incubation of both enzymes with their most probable substrates (L-histidine, NADH and αKG) did not lead to the formation of an additional radiolabelled product as for the case of staphylopine biosynthesis (17) (Figure 3C). One possibility was therefore that the product of PaCntM was migrating at the same position as the yNA in the conditions used during the TLC separation. We therefore decided to study the same co-incubations by HILIC/ESI-MS, following the mass expected for the yNA intermediate and the pseudopaline found in the extracellular fraction of *P. aeruginosa* grown in MS medium. These experiments confirmed that the incubation of PaCntL with SAM and L-histidine led to the formation of the yNA reaction intermediate (Figure 3D, top), and most of all, revealed the production of pseudopaline when co-incubating PaCntL, PaCntM and their proposed substrates (SAM, L-histidine, NADH and αKG; Figure 3D, bottom). Co-incubations using alternative substrates of PaCntM (pyruvate or NADPH) only led to the production of yNA. Interestingly, pseudopaline and yNA eluted at the same volume in these HILIC-ESI/MS experiments, showing that their physical properties are very similar, as suggested by our previous TLC experiments.

**Figure 3:**
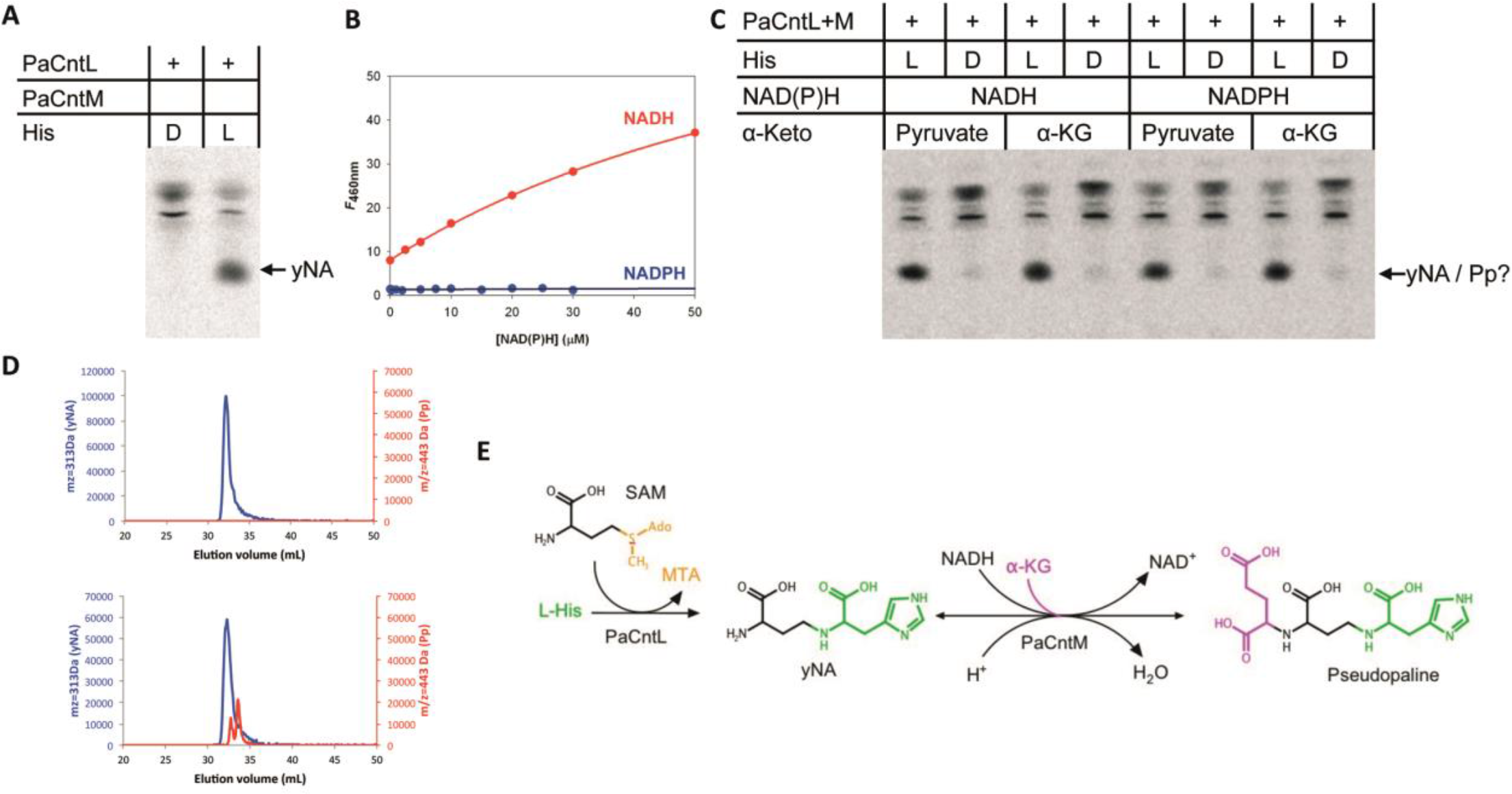
*In vitro* reconstitution of the pseudopaline biosynthesis pathway. (A) TLC experiment using PaCntL and [^14^C]-SAM showing that PaCntL discriminates between D- and L-histidine substrate with the production of the reaction intermediate (noted yNA) only visible when using L-histidine. (B) Titration of NADPH (blue) and NADH (red) binding to PaCntM (5µM) followed by fluorescence resonance energy transfer. Fitting of the data obtained for NADH led to a K_d_ of 30µM. (C) TLC separation of reaction products incubating [^14^C]-SAM using purified enzymes (PaCntL and PaCntM), different source of α-ketoacid (pyruvate or α-KG), cofactor (NADH or NADPH) and histidine (L-His or D-His). (D) HILIC/ESI-MS chromatograms of putative reaction products using PaCntL incubated with L-histidine, revealing the production of the yNA intermediate (top), and a mix of PaCntL and PaCntM incubated with all their putative substrate (SAM, L-histidine, NADH and α-Ketaoglutarate), revealing the specific detection of pseudopaline in this case (red trace, bottom). (E) Summary of the PaCntL/M-dependent biosynthesis pathway for the assembly of pseudopaline from L-his, SAM, NADH and α-KG.

Pseudopaline is therefore biosynthesized in two steps: first, a nucleophylic attack of one αaminobutyric acid moiety from SAM onto L-histidine catalysed by PaCntL to produce the reaction intermediate yNA, and second, a NADH reductive condensation of the yNA intermediate with a molecule of αKG catalysed by PaCntM to produce pseudopaline (Figure 3E). Pseudopaline differs from staphylopine by the stereochemistry of the histidine moiety (L- and D- respectively) and by the presence of an αKG moiety instead of pyruvate in staphylopine. The biosynthesis of a specific metallophore by different bacteria recalls the chemical evolution of a large diversity of siderophore in a chemical rivalry to get access to one’s own pool of metal (28). Indeed, once in the extracellular medium, secreted metallophores are a common good, and a privileged access presumably gives a selective advantage.

### Pseudopaline is involved in nickel and zinc uptake, depending on the chelating properties of the media

In order to address the involvement of pseudopaline in metal uptake *in vivo*, we compared the intracellular concentration of various metals in PA14 WT, *ΔcntL* and *ΔcntL::cntL* strains. Cells were grown in pseudopaline-synthesis conditions determined above (MS medium) and the intracellular metal concentration was measured by ICP-MS. Under these growth conditions we observed a significant 90% reduction of intracellular nickel concentration in the *ΔcntL* mutant strain (Figure 4A), which was mostly recovered in the complemented strain. The levels of all the other metals were not changed in the *ΔcntL* mutant strain compared to the WT strain (data not shown). A similar 90% reduction in intracellular nickel concentration was also observed when the culture was supplemented with 1μM NiCl_2_ (Figure S4), thus confirming that nickel uptake was predominantly performed by pseudopaline in these metal-poor media. We were intrigued by the apparent contradiction between the clear *cnt* operon regulation by zinc, and the absence of any effect on zinc uptake. A possible explanation is that the effect of *cnt* could be masked by the effect of a zinc ion importer such as the ZnuABC zinc transport system described in *P. aeruginosa* (22). In an attempt to discriminate between both transport systems, we sequestered free metal ions by supplementing the growth medium with increasing concentrations of EDTA, a chelating agent for divalent metals. Interestingly, although we did not observe any effect using 10 μM EDTA, the supplementation with 100μM EDTA ultimately revealed a pseudopaline-dependent zinc uptake, with a 60% decrease of intracellular zinc content in the *ΔcntL* mutant strain in comparison with the WT strain (Figure 4B). The complemented strain accumulated zinc to a level comparable to the WT. In these chelating conditions the pseudopaline-dependent nickel import is abolished (Figure 4A), hence proving a direct link between pseudopaline and zinc uptake in metal scarce conditions with competing zinc chelators. These conditions may prevail at the host-pathogen interface where metal binding proteins such as calprotectin are produced by the host (29, 30), or in AMS where metals are complexed in a nutritional immunity framework (1, 21).

**Figure 4:**
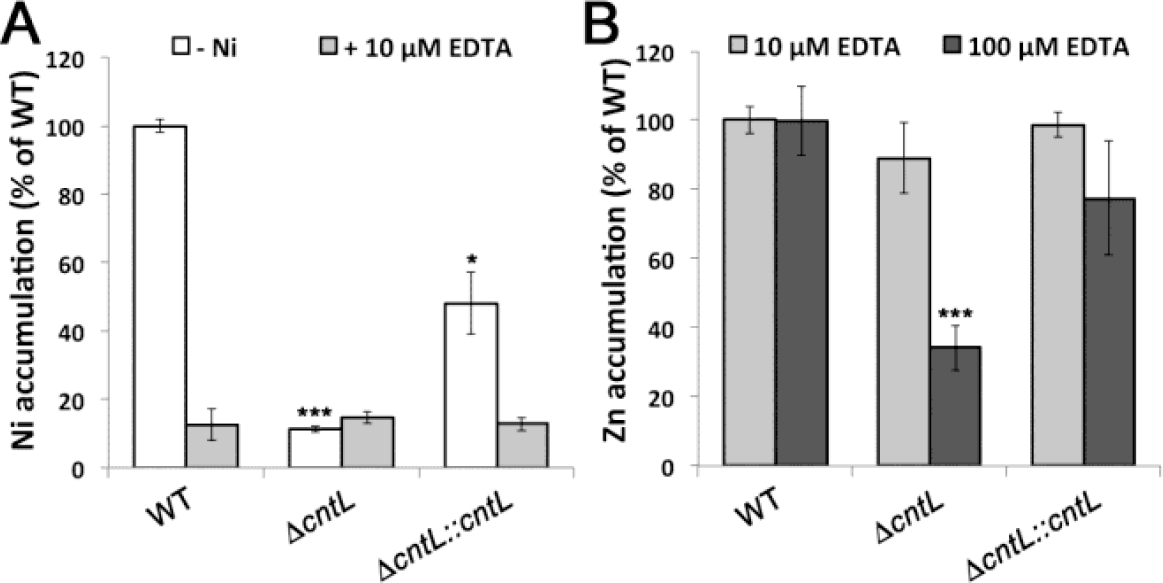
Pseudopaline is involved in nickel uptake in minimal media and in zinc uptake in chelating media. Intracellular nickel (A) or zinc (B) levels measured by ICP-MS in WT, *ΔcntL* and *ΔcntL::cntL* strains grown in MS medium supplemented or not with 10 or 100µM EDTA. Error bars, mean ± s.d. \**P*<0.05, \*\**P*<0.01 and \*\*\**P*<0.001 as compared to the WT.

### Model of pseudopaline synthesis and transport pathway in *P. aeruginosa*

We next investigated the putative roles of the two membrane proteins that are found in the *cnt* operon of *P. aeruginosa* by determining the pseudopaline level in the extracellular and intracellular fractions of WT and mutant strains (Figure 5A and 5B, respectively). With regard to PaCntO, we found a small decrease in the extracellular content of pseudopaline in the Δ*cntO* mutant strain in comparison with the WT strain. However, we also found that this Δ*cntO* mutant strain was partly impaired in nickel accumulation (Figure S5). Altogether, and because PaCntO belongs to the TBDT family of extracellular transporter, its most probable role is in the import of pseudopaline-metal complexes, although it is not excluded that other proteins of this family could participate in this process. Next, we noted a large decrease in the extracellular pseudopaline level in the Δ*cntI* mutant strain in comparison with the WT strain, with a concomitant increase in the intracellular space, consistent with a role of PaCntI in pseudopaline export. It is also interesting to note that a Δ*cntI* mutant strain is virtually unable to grow in AMS (21). The most probable scenario is that this mutant is deficient in metal content, including zinc, but pseudopaline accumulation in the cytoplasmic space actually worsens the situation by chelating an already poorly available metal. This assumption is supported by our finding that a double Δ*cntL*Δ*cntI* mutant supresses the detrimental growth defect of the single Δ*cntI* mutant strain, *ie* the absence of pseudopaline restores the normal growth of a mutant devoid of the pseudopaline exporter (Figure S6). A model recapitulating the pseudopaline pathway is shown in Figure 5.

**Figure 5:**
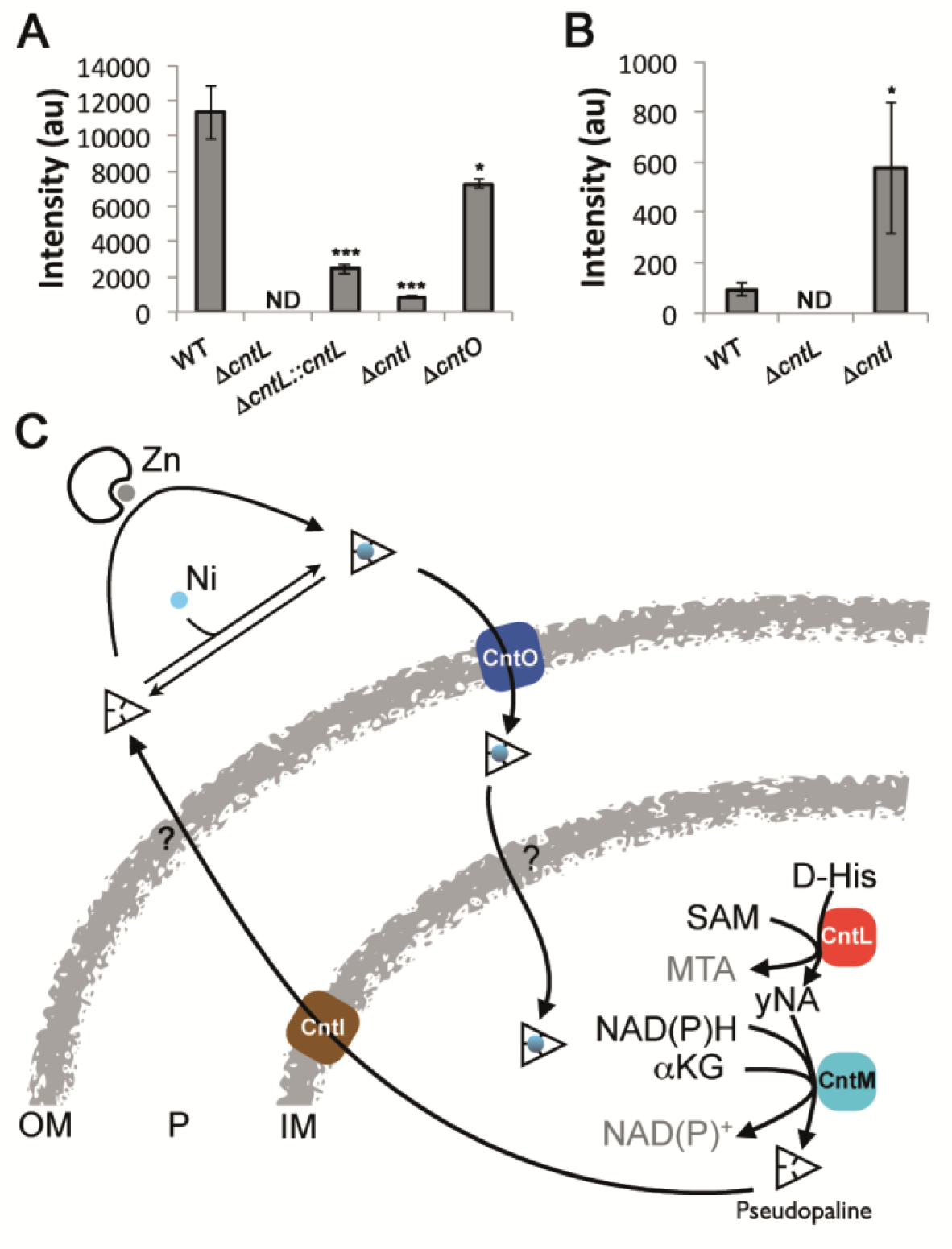
Model of pseudopaline synthesis, secretion and metal uptake in *P. aeruginosa*. (A) Extracellular detection of pseudopaline in the extracellular fraction of WT and mutant strains. Error bars, mean ± s.d. \**P*<0.05, \*\**P*<0.01 and \*\*\**P*<0.001 as compared to the WT. (B) Intracellular detection of pseudopaline in the intracellular fraction of WT and mutant strains. ND: Not Detectable. (C) Model of pseudopaline production, secretion and recovery of nickel or zinc. Outer membrane (OM), inner membrane (IM), periplasm (P).

It is interesting to note the differences and similarities between staphylopine and pseudopaline and between their respective biosynthetic pathways (Figure S7). On one hand, pseudopaline differs from staphylopine by the incorporation of a L-histidine instead of a D-histidine moiety in staphylopine, thus explaining the absence of amino acid racemase in *P. aeruginosa*. Another particularity of pseudopaline is the use of an αKG moiety instead of pyruvate as substrate for the second reaction mediated by PaCntM. Together this leads to two species-specific metallophores that might give a selective advantage in a competing environment. The fact that staphylopine and pseudopaline belong to Gram-positive and Gram-negative bacteria has important consequences on their respective transport mechanisms across the two types of bacterial envelopes. Although the transporters of staphylopine are well identified, the outer membrane exporter pseudopaline and inner membrane importer of the pseudopaline-metal complex remains to be discovered (Figure 5). Recycling of the metallophore could also take place in *P. aeruginosa*, as recently exemplified in the case for pyoverdine (31). An interesting aspect of this work is the discovery of two different pathways for the export of these nicotianamine-like bacterial metallophores. Whereas *S. aureus* uses a protein belonging to the MFS family (SaCntE) for staphylopine export, *P. aeruginosa* uses a protein belonging to the DMT family of transporters, with PaCntI possessing two predicted EamA domains for pseudopaline export. In the view of their importance in the growth or virulence of these important human pathogens (19, 21), they could emerge as attractive targets for novel antibiotic development.

## ACKNOWLEDGEMENTS

This work was supported by the ANR-14-CE09-0007-03 grant allocated to PA, RV, RL and EBD and the grant from “Vaincre la Mucoviscidose” (RFI20160501495) allocated to RV and PA. We thank B. Douzi for fruitful discussions, B. Ize for RNA preparation and C. Soscia for technical support as well as O. Uderso, I. Bringer and A. Brun for material and media preparations.

